# Whole Genome Sequencing of *Plasmodium vivax* Isolates Reveals Frequent Sequence and Structural Polymorphisms in Erythrocyte Binding Genes

**DOI:** 10.1101/2020.03.23.003293

**Authors:** Anthony Ford, Daniel Kepple, Beka Raya Abagero, Jordan Connors, Richard Pearson, Sarah Auburn, Sisay Getachew, Colby Ford, Karthigayan Gunalan, Louis H. Miller, Daniel A. Janies, Julian C. Rayner, Guiyun Yan, Delenasaw Yewhalaw, Eugenia Lo

## Abstract

*Plasmodium vivax* malaria is much less common in Africa than the rest of the world because the parasite relies primarily on the Duffy antigen/chemokine receptor (*DARC*) to invade human erythrocytes, and the majority of Africans are Duffy negative. Recently, there has been a dramatic increase in the reporting of *P. vivax* cases in Africa, with a high number of them being in Duffy negative individuals, potentially indicating *P. vivax* has evolved an alternative invasion mechanism that can overcome Duffy negativity. Here, we analyzed single nucleotide polymorphism (SNP) and copy number variation (CNV) in Whole Genome Sequence (WGS) data from 44 *P. vivax* samples isolated from symptomatic malaria patients in southwestern Ethiopia, where both Duffy positive and Duffy negative individuals are found. A total of 236,351 SNPs were detected, of which 21.9% was nonsynonymous and 78.1% was synonymous mutations. The largest number of SNPs were detected on chromosomes 9 (33,478 SNPs; 14% of total) and 10 (28,133 SNPs; 11.9%). There were particularly high levels of polymorphism in erythrocyte binding gene candidates including reticulocyte binding protein 2c (*RBP*2c), merozoite surface protein 1 (*MSP*1), and merozoite surface protein 3 (*MSP*3.5, *MSP*3.85 and *MSP*3.9). Thirteen genes related to immunogenicity and erythrocyte binding function were detected with significant signals of positive selection. Variation in gene copy number was also concentrated in genes involved in host-parasite interactions, including the expansion of the Duffy binding protein gene (*PvDBP*) on chromosome 6 and several *PIR* genes. Based on the phylogeny constructed from the whole genome sequences, the expansion of these genes was an independent process among the *P. vivax* lineages in Ethiopia. We further inferred transmission patterns of *P. vivax* infections among study sites and showed various levels of gene flow at a small geographical scale. The genomic features of *P. vivax* provided baseline data for future comparison with those in Duffy-negative individuals, and allowed us to develop a panel of informative Single Nucleotide Polymorphic markers diagnostic at a micro-geographical scale.

## Introduction

Vivax malaria is the most geographically widespread human malaria, causing over 130 million clinical cases per year worldwide [1]. *Plasmodium vivax* can produce dormant liver-stage hypnozoites within infected hosts, giving rise to relapse infections from months to years. This unique feature of *P. vivax* contributes to an increase in transmission potential and increases the challenge of elimination [2]. Understanding *P. vivax* genome variation will advance our knowledge of parasite biology and host-parasite interactions, as well as identify potential drug resistance mechanisms [3, 4]. Such data will also help identify molecular targets for vaccine development [5-7], and provide new means to track the transmission and spread of drug resistant parasites [8-9].

Compared to *P. falciparum, P. vivax* isolates from Southeast Asia (e.g., Thailand and Myanmar), Pacific Oceania (Papua New Guinea), and South America (Mexico, Peru, and Colombia) have significantly higher nucleotide diversity at the genome level [2]. This could be the result of frequent gene flow via human movement, intense transmission, and/or variation in host susceptibility [10-14]. *P. vivax* infections are also much more likely to contain multiple parasite strains in areas where transmission is intense and/or relapse is common [10, 15-18]. In Papua New Guinea, for example, *P. vivax* infections had an approximately 3.5-fold higher rate of polyclonality and nearly double the multiplicity of infection (MOI) than the *P.* falciparum infections [16]. Similar rates of polyclonality and MOI have also been reported in *P. vivax* in Cambodia [6]. It is possible intense transmission has sustained a large and stable parasite population in these regions [17,18]. By contrast, geographical differentiation and selection pressure over generations can lead to fixation of parasite genotypes in local populations. In the Asia-Pacific region, *P. vivax* showed a high level of genetic relatedness through inbreeding among the dominant clones, in addition to strong selection imposed in a number of antimalarial drug resistance genes [19]. In Ethiopia, the chloroquine resistance transporter gene (*Pvcrt*) of *P. vivax* on chromosome 14 had been shown with significant selection in a region upstream of the promotor, highlighting the ability of *P. vivax* to rapidly evolve in response to control measures [20]. Apart from mutations, high copy number observed in *Pvcrt* and multidrug resistant gene (*Pvmdr*1) has also been shown to be associated with increased antimalaria drug resistance [21,22].

Recent genomic studies have indicated that some highly polymorphic genes in the *P. vivax* genome are associated with red blood cell invasion and immune evasion [10, 12, 19, 23]. They include the merozoite surface protein genes *MSP*1 (PVP01_0728900) and *MSP*7 (PVX_082665), Pv-fam-b (PVX_002525), Pv-fam-e (PVX_089875), the reticulocyte binding protein gene *RBP*2c (PVP01_0534300), serine-repeat antigen 3 (*SERA*; PVX_003840), as well as virulent genes (*VIR*) such as *VIR*22 (PVX_097530) and *VIR*12 (PVX_083590) [23-29]. Polymorphisms in genes associated with immune evasion and reticulocyte invasion have important implications for the invasion efficiency and severity of *P. vivax* infections. Members of the erythrocyte binding gene family, including reticulocyte binding proteins (*RBP*s), Duffy-binding proteins (*DBP*s), and merozoite surface proteins (*MSP*3 and *MSP*7) have been previously shown to exhibit high sequence variation in *P. vivax* [20, 30]. The polymorphisms in *RBP*1 and *RBP*2 genes may relate to an increased capability of erythrocyte invasion by *P. vivax* [31-33]. It has been suggested that Pv*RBP*2b-TfR1 interaction is vital for the initial recognition and invasion of host reticulocytes [34], prior to the engagement of *PvDBP1* and Duffy antigen chemokine receptor (*DARC*) and formation of a tight junction between parasite and erythrocyte [35]. Apart from Pv*RBP*, Reticulocyte Binding Surface Antigen (Pv*RBSA*) [36], an antigenic adhesin, may also play a key role in *P. vivax* parasites binding to target cells, possessing the capability of binding to a population of reticulocytes with a different Duffy phenotype [37, 38]. Another erythrocyte binding protein gene (Pv*EBP*), a paralog of *PvDBP1*, which harbors all the hallmarks of a *Plasmodium* red blood cell invasion protein, including conserved Duffy-binding like and C-terminal cysteine-rich domains [39], has been recently shown to be variable in copy number in the Malagasy *P.* vivax [39]. Functional analyses indicated that region II of this gene bound to both Duffy-positive and Duffy-negative reticulocytes, although at a lower frequency compared to *PvDBP*, suggestive of its role in erythrocyte invasion [40]. Both Pv*EBP*1 and Pv*EBP*2 genes exhibit high genetic diversity and are common antibody binding targets associated with clinical protection [41, 42]. Other proteins such as tryptophan-rich antigen gene (*TRAg*), anchored micronemal antigen (*GAMA*), and Rhoptry neck protein (*RON*) have also been suggested to play a role in red cell invasion, especially in low-density infections [43-47]. Information of the polymorphisms in these genes will have important implications on the dynamics of host-parasite interactions.

Compared to Southeast Asia and South America where *P. vivax* is highly endemic, data on polymorphisms in erythrocyte binding gene candidates of *P. vivax* from Africa is limited. Filling the gap is critical for identifying functional genes in erythrocyte invasion, biomarkers for tracking the African *P. vivax* isolates, as well as potential gene targets for vaccine development. It was previously thought that most African populations were immune to *P.* vivax infections due to the absence of *DARC* gene expression required for erythrocyte invasion. However, several recent reports have indicated the emergence and potential spread of *P. vivax* across Africa [32, 48-50]. The objective of this study was to describe genomic variation of *P. vivax* from Ethiopia. Specifically, we examined the level of genetic polymorphisms in a panel of 64 potential erythrocyte binding protein genes that have been suggested to play a role in the parasite-host invasion process. In addition, we inferred transmission patterns of *P. vivax* infections from different study sites based on the genetic variants. A recent study by Auburn *et al*. [20] has compared the genetic variants of *P. vivax* from Ethiopia with other geographical isolates. In the present study, we focus on the genomic characteristics of *P. vivax* among different study sites in Ethiopia with the goals to establish a baseline for genome comparison with the Duffy-negative *P. vivax* in our ongoing investigation, as well as to develop a panel of informative Single Nucleotide Polymorphic (SNP) markers diagnostic at a micro-geographical scale.

## Materials and Methods

### Ethics statement

Scientific and ethical clearance was obtained from the Institutional Scientific and Ethical Review Boards of Jimma and Addis Ababa Universities in Ethiopia, and The University of North Carolina, Charlotte, USA. Written informed consent/assent for study participation was obtained from all consenting heads of households, parents/guardians (for minors under age of 18), and each individual who was willing to participate in the study.

### Study area and sample collection

Genomic DNA was extracted from 22 clinical samples collected in Jimma, southwestern Ethiopia during peak transmission season (September – November, 2016; Figure 1). Finger-pricked blood samples were collected from malaria symptomatic (who has fever with axillary body temperature > 37.5°C and with confirmed asexual stages of malaria parasite based on microscopy) or febrile patients visiting the health centers or hospitals at each of the study sites. Thick and thin blood smears were prepared for microscopic examination, and 4-6 ml of venous blood were collected from each *P. vivax*-confirmed patient in K2 EDTA blood collection tubes. For the whole blood samples, we used the Lymphoprep/Plasmodpur-based protocol to deplete the white blood cells and enrich the red blood cell pellets [51]. DNA was then extracted from approximately 1 ml of the red blood cell pellets using Zymo Bead Genomic DNA kit (Zymo Research) following the manufacturer’s procedures. The extracted DNA were first assessed by nested and quantitative PCR methods to confirm and quantify *P. vivax* of the infected samples [52]. From a larger set of samples, we then performed microsatellite analyses using seven different loci [53]. Only monoclonal samples were selected and proceeded for sequencing. Whole genome sequencing was conducted on the Illumina HiSeq 3000 Sequencing Platform at the Wellcome Sanger Institute (European Nucleotide Archive [ENA] accession number of each sample in Table 1). The generated sequence reads were mapped individually to the publicly available reference genome PvP01 from Gene DB using Bowtie version 2 [54]. The original 22 samples were processed to remove reads other than *P. vivax*. The percentage coverage of the *P. vivax* reads in our samples were high enough to not affect the results. An additional 24 sample sequence data were obtained as FASTQ files from the ENA. These samples were collected from Arbaminch, Badowacho, Halaba, and Hawassa in southwestern Ethiopia (Figure 1), the Duffy status of each of these 24 samples is unknown. They were then aligned to the PVP01 reference genome using BWA-MEMv.2 with default settings [55, 56]. The overall quality of each resulting BAM was assessed using FASTQC. Similarly, we concluded that the percentage of the *P. vivax* reads covered in the additional 24 samples were high enough to reflect the dominant signal of the variants and negate polyclonal influences. Two of our samples displayed a significant decline in average quality in read mapping and were therefore removed from further SNP variant and copy number variation analyses.

**Figure 1.**
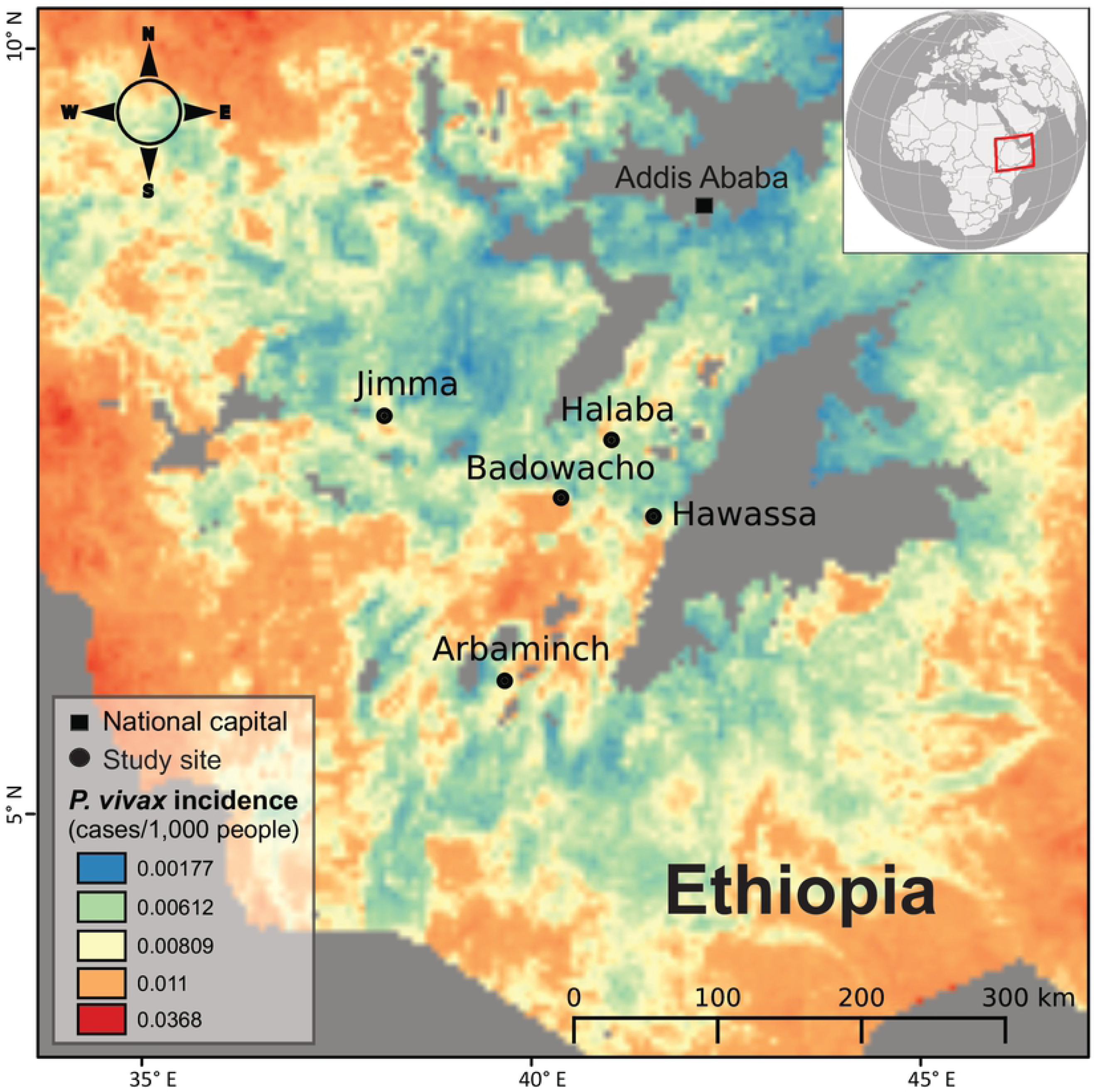
An overview of the *P vivax* sample collection locations including Arbaminch, Badowacho, Hawassa, Halaba, and Jimma in southwestern Ethiopia.

### SNP discovery, annotation, and filtering

Potential SNPs were identified by SAM tools v.1.6 mpileup procedure [57] in conjunction with BCF tools v.1.6 [57] across all 44 sample BAM files using the PVP01 reference genome. Compared to the Salvador-I, the PVP01 reference genome consists of 14 major chromosomal sequences, and provides a greater level of gene annotation power and improved assembly of the subtelomeres [56]. We analyzed only sequence reads that were mapped to these 14 major chromosomal sequences. The hypervariable and subtelomeric regions in our samples were retained during the variant calling procedure and each sample BAM file had duplicates marked using SAMtools 1.6 markdup procedure. For the mpileup procedure, the maximum depth threshold, which determines the number of maximum reads per file at a position, was set to 3,000 million to ensure that the maximum amount of reads for each position was not reached. Samples were pooled together using a multisampling variant calling approach. The SNPs were then annotated with SnpEff v.4.3T [58] based on the annotated gene information in GeneDB. Filtering was done using the following standard metrics, including Read Position Bias, Mapping Quality vs Strand Bias, Raw read depth, Mapping Quality Bias, Base Quality Bias, and Variant Distant Bias produced by SAM tools and BCF tools during the variant calling procedure. In Snp Sift, data was filtered by choosing SNPs that had a Phred Quality score ≥40, a raw read depth (DP) ≥30, and a base quality bias >0.1 [59]. We then calculated the allele frequency for each SNP position for all 44 samples using the frequency procedure in VCF tools v.0.1.15 [60]. The total number of SNPs across all samples, as well as the number of nonsynonymous and synonymous mutations were recorded. Mutations were compared among the 14 chromosomes in addition to a panel of 64 erythrocyte binding genes.

### Copy number variation analyses

Copy number variation of gene regions was assessed with CNVnator [61]. CNVnator uses mean-shift theory, a partitioning procedure based on an image processing technique and additional refinements including multiple bandwidth partitioning and GC correction [61]. We first calculated the read depth for each bin and correct GC-bias. This was followed by mean-shift based segment partition and signal merging, which employed an image processing technique. We then performed CNV calling, of which segments with a mean RD signal deviating by at least a quarter from genomic average read depth signal were selected and regions with a *P*-value less than 0.05 were called. A one-sided test was then performed to call additional copy number variants. SAM tools v.1.6 was utilized in our data preprocessing step to mark potential duplicates in the BAM files and followed the CNV detection pipeline [62]. We extracted the read mappings from each of BAM files for all chromosomes. Once the root file was constructed using the extracted reads, we generated histograms of the read depths using a bin size of 100. The statistical significance for the windows that showed unusual read depth was calculated and the chromosomes were partitioned into long regions that have similar read depth.

To validate the results from CNVnator, we used the GATK4 copy number detection pipeline to further examine gene copy number [63-65]. The read coverage counts were first obtained from pre-processed genomic intervals of a 1000-bp window length based on the PvP01 reference genome. The read fragment counts were then standardized using the Denoise Read Counts that involved two transformations. The first transformation was based on median counts, including the log_2_ transformation, and the counts were normalized to center around one. In the second transformation, the tool denoises was used to standardized copy ratios using principal component analysis.

### Test for positive selection

Regions of positive selection were examined among the 44 Ethiopian *P. vivax* isolates using the integrated haplotype score approach, specifically the SciKit-Allel for python, a package used for analysis of large scale genetic variation data [66]. Before the samples were run through Scikit-Allel, genotypes for each of the samples were phased using BEAGLE [67]. Genes that were detected with signals of positive selection by SciKit-Allel, as well as a panel of 64 potential erythrocyte binding genes were further evaluated using the PAML package (Phylogenetic Analysis by Maximum Likelihood) [68]. Using the codeml procedure in PAML, DNA sequences were analyzed with the maximum likelihood approach in a phylogenetic framework. The synonymous and nonsynonymous mutation rates between protein-coding DNA sequences were then estimated in order to identify potential regions of positive selection. We created two models, the neutral model M1 and the selection model M2. The average d_N_/d_S_ values were estimated across all branches in both M1 and M2 models and the average d_N_/d_S_ values across all sites in the M2 model. The d_N_/d_S_ values were compared between the two models using a likelihood ratio test for significant positive selection.

### Comparison of nucleotide diversity among EBP gene regions

Based on the literature [23-33], we identified 64 gene regions that are potentially related to erythrocyte binding in *P. vivax* (Supplementary Table 1). These included the *DBP* (duffy binding protein), *EBP* (erythrocyte binding protein), *MSP* (merozoite surface protein), and *RBP* (reticulocyte binding protein) multigene families, the tryptophan rich antigen gene family (*TRAg*), GPI-anchored microanemal antigen (*GAMA)*, microneme associated antigen (*MA)*, rhoptry associated adhesin (*RA)*, high molecular weight rhoptry protein 3 (*RHOP*3), and rhoptry neck protein (*RON)* genes. Previous study has shown that the transcriptome profiles of the *TRAg* genes were differentially transcribed at the erythrocytic stages, indicating that these genes may play specific roles in blood-stage development [43]. The reticulocyte binding protein multigene family encodes genes that each have a receptor on the surface that is essential for the host-invasion stage of *P. vivax* [69]. The *MSP* multigene family, currently assumed to be a candidate for vaccine generation, also plays a role in the invasion stage of *P. vivax* and is also immunogenic [26]. The nucleotide diversity of 64 potential erythrocyte binding genes were compared among the 44 *P. vivax* sample consensus sequences using DnaSP [70]. The Pairwise-Deletion method where gaps were ignored in each pairwise comparison was used for this calculation.

### Genetic relatedness and transmission network analyses

Phylogenetic analyses were performed to infer the genetic relatedness among the 44 Ethiopian isolates. Sequence alignment was first conducted using a multiple sequence alignment program in MAFFT v. 7 [71]. The alignment was then trimmed to remove gaps using trimal (the *gappyout* option) that trimmed the alignments based on the gap percentage count over the whole alignment. After sequence editing, we concatenated all alignment files using FASconCAT-G [72], a perl program that allows for concatenation and translation (nucleotide to amino acid states) of multiple alignment files for phylogenetic analysis. We used the maximum likelihood method implemented in the Randomized Accelerated Maximum Likelihood (RAxML) v8 to construct phylogenetic trees [73]. The GTRGAMMA model was used for the best-scoring maximum likelihood tree. The GTR model incorporates the optimization of substitution rates and the GAMMA model accounts for rate heterogeneity. A total of 100 rapid bootstrap runs were conducted to evaluate the confidence of genetic relationships. In addition, we performed principal component analyses using the glPCA function in R, a subset of the adegenet package [74], to determine the genetic relatedness of the samples among the different study sites in Ethiopia. A transmission network was created using StrainHub, a tool for generating transmission networks using phylogenetic information along with isolate metadata [75]. The transmission network was generated using the locations of the samples as the nodes and calculating the source hub ratio between each location. The source hub ratio was calculated by the number of transitions originating from a node over the total number of transitions related to that node. A node with a ratio close to 1 indicates a source, a ratio close to 0.5 indicates a hub, and a ratio close to 0 indicates a sink for the *P. vivax* infections.

## Results

### Distribution of SNPs among the chromosomes and EBP genes

A total of 252,973 SNPs were detected among the 44 Ethiopian *P. vivax* samples (Figure 2), with 21.5% (54,336 out of 252,973) nonsynonymous and 78.5% (198,637 out of 252,973) synonymous mutations (Figure 3A). The highest number of SNPs were observed on chromosomes 7 (28,856 SNPs; 11.4%), 9 (28,308 SNPs; 11.2%), and 12 (28,190 SNPs; 11.1%); whereas the lowest number of SNPs were observed on chromosomes 3 (6,803 SNPs; 2.7%), 6 (5,044 SNPs; ∼2%), and 13 (8,809 SNPs; 3.5%; Figure 3A; Supplementary Table 2).

**Figure 2.**
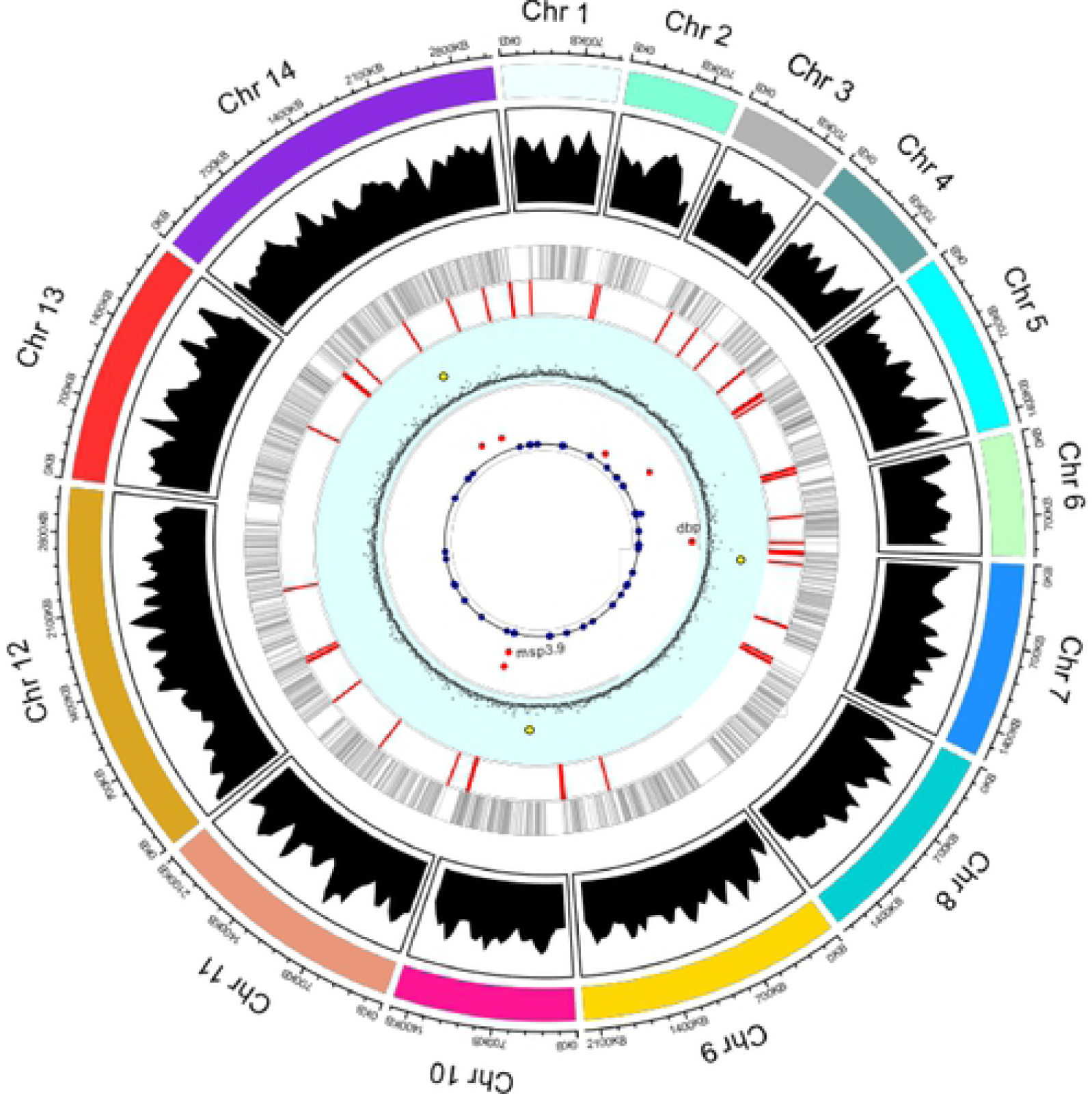
A summary representation of the *P. vivax* genome, with the outer ring as an ideogram representing the 14 nuclear chromosomes and sizes of each. The second track represented the average coverage for each chromosome among the 44 Ethiopian samples. The third track containing the gray vertical dashes represented the distribution of genes across the 14 chromosomes. The forth track that contained the red vertical lines represented the 64 erythrocyte binding gene candidates. The fifth inner track with the light blue background represented the d_N_/d_S_ ratio calculated by partitioning the chromosomes into genomic regions and d_N_/d_S_ directly. The three outliers (yellow dots) represented three unknown plasmodium protein genes that were detected with significant positive selection. The sixth track indicated the overall copy number variation calculated using CNVnator. Red dots represented genes with copy number variation among the Ethiopian genomes.

**Figure 3.**
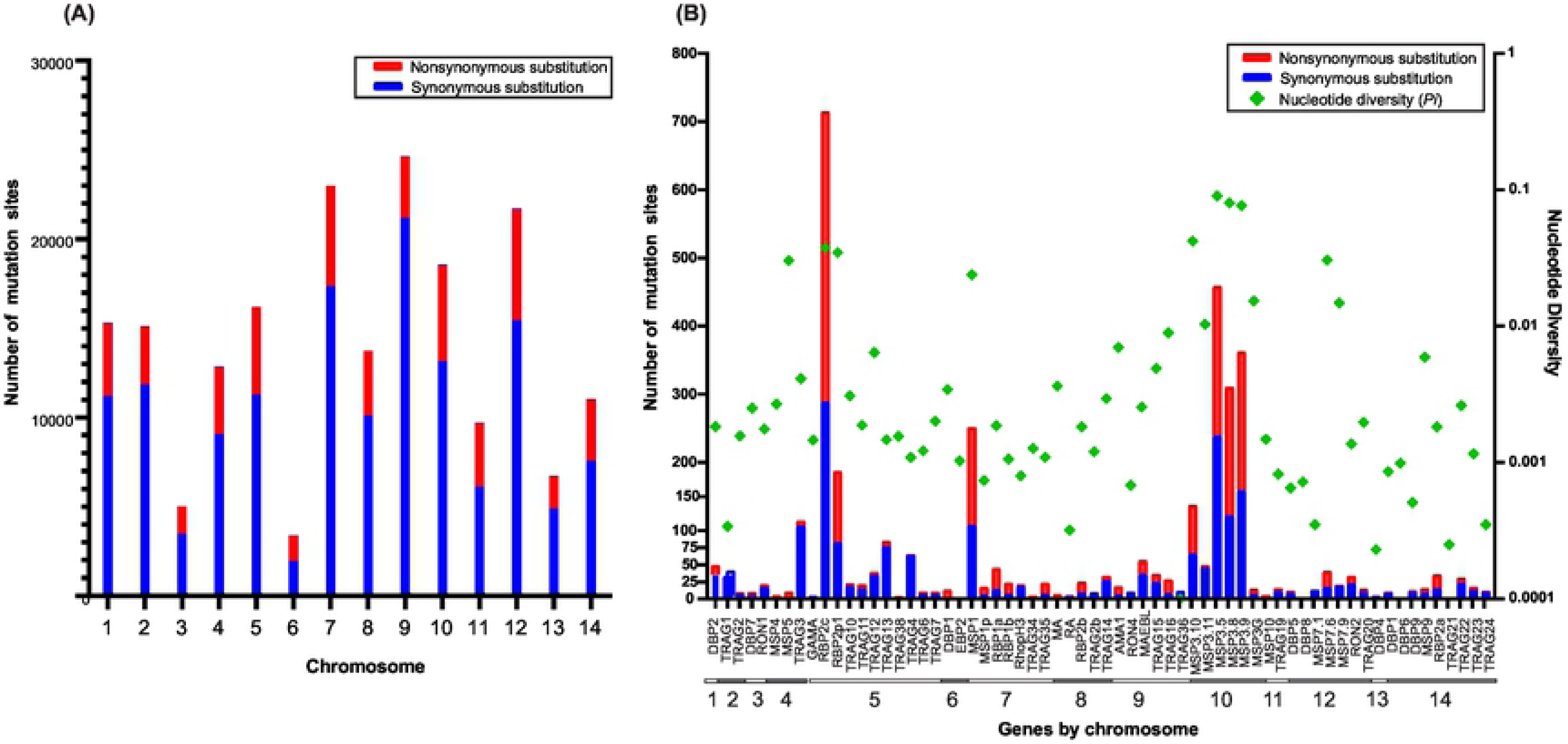
(A) A distribution of the nonsynonymous and synonymous mutations of each chromosome. A higher proportion of synonymous mutations was observed compared to nonsynonymous mutations. Chromosomes 7, 9, and 12 have the most mutations overall, with chromosomes 6 and 3 having the fewest number of mutations. (B) Number of mutation sites and the nucleotide diversity of 64 erythrocyte binding genes. The *PvRBP* and *PvMSP* multigene families have the highest number of polymorphic sites when compared to the others, with *PvRBP*2c the highest number of nonsynonymous and synonymous mutations, followed by *PvMSP*3 and *PvMSP*1. Approximately 40% of the mutations were nonsynonymous. These genes were also indicated with the highest nucleotide diversity.

The 64 erythrocyte binding genes accounted for 3,607 of the total SNPs, with 1685 (46.7%) identified as nonsynonymous and 1922 (53.3%) as synonymous mutations (Figure 3B). Among these genes, the highest number of SNPs were observed in reticulocyte binding protein gene (*RBP*2c) on chromosome 5, followed by the *MSP*3 multigene family (*MSP*3.5, *MSP*3.9 and *MSP*3.8) on chromosome 10. Nucleotide diversity also showed to be highest in the *RBP* and *MSP*3 multigene families, with an average nucleotide diversity of 1.3% and 2.8%, respectively, among our samples (Figure 3B). By contrast, the lowest number of SNPs were observed in the Duffy binding protein gene (*DBP*1) on chromosome 6 with a total of 13 SNPs, of which 12 were identified as nonsynonymous and one as synonymous mutations (Figure 3B). Likewise, another erythrocyte binding protein (*EBP*2), located also on chromosome 6, was one of the least variable genes with only one nonsynonymous mutation. The *TRAg* gene family also showed a low level of nucleotide diversity when compared to the other *EBP* gene families with an average nucleotide diversity of 0.2% (Figure 3B).

### Gene regions under positive selection

Based on the integrated haplotype scores, positive selection was detected in 13 gene regions (Figure 4). These included the sub-telomeric protein 1 (*STP*1) on chromosome 5, the membrane associated erythrocyte binding-like protein (*MAEBL*) on chromosome 9, *MSP*3.8 on chromosome 10, as well as various plasmodium interspersed repeats (*PIR)* protein genes on chromosomes 3, 5, 7, 10, 11, and 12 (Figure 4). Based on PAML, 25 out of the 64 erythrocyte binding genes showed evidence of positive selection (Table 2; Supplementary Table 3). The majority of these genes belong to the *TRAg* multigene family. The *TRAg* genes had an average d_N_/d_S_ ratio of 2.75 across all branches and an average of 5.75 across all sites for the M2 model tested for selection (Table 2). Compared to the other *TRAg* genes, *TRAg*15 had more sites detected under positive selection, with 50 of the sites showing a posterior probability greater than 50% and 43 showing a posterior probability greater than 95% (Table 2). While the *TRAg*4 gene had the highest d_N_/d_S_ ratio across all sites among other *TRAg* genes, only six sites were shown under positive selection with a posterior probability greater than 50% and one with a posterior probability greater than 95%.

**Figure 4.**
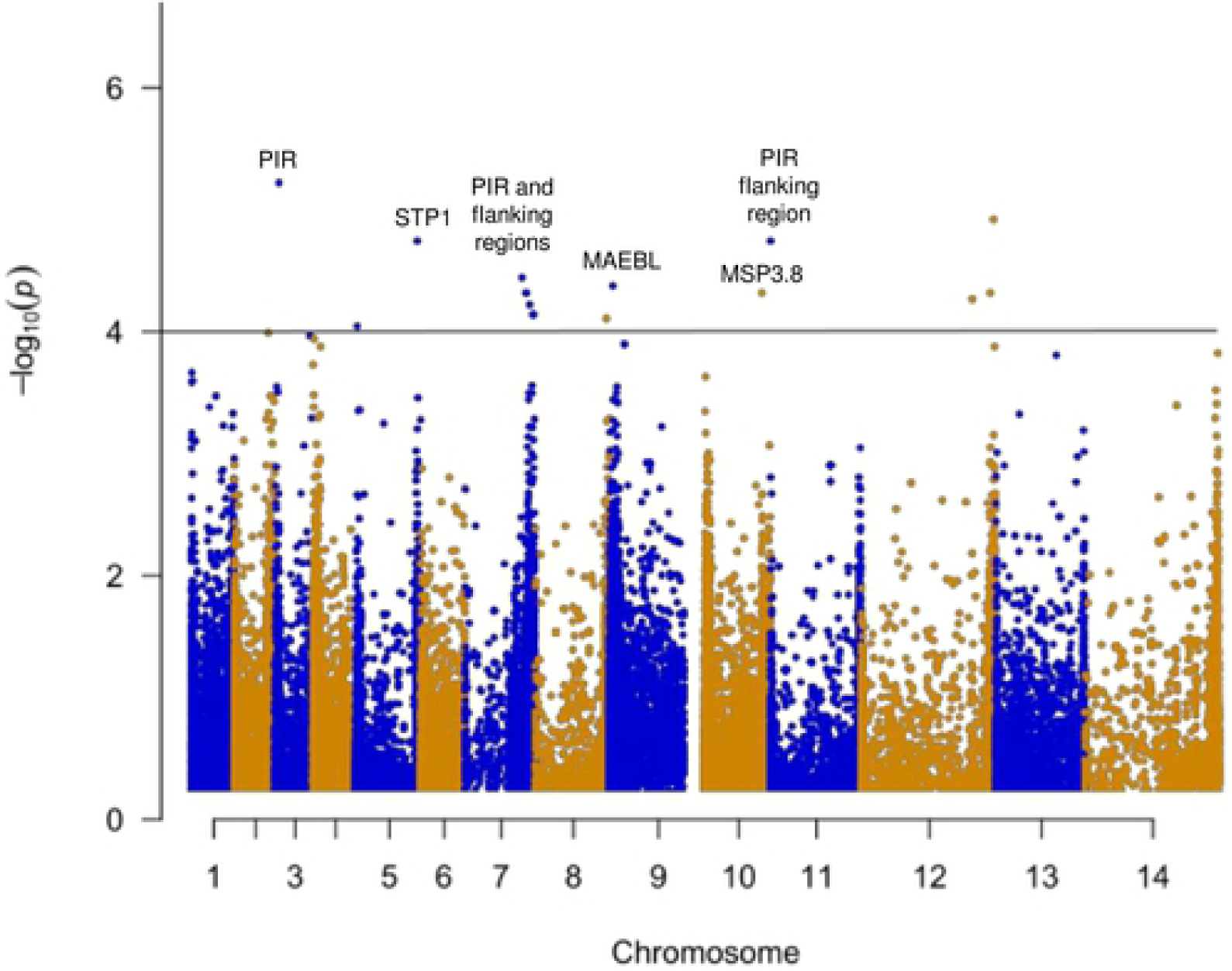
Signal of positive selection across the 14 chromosomes among all *P. vivax* samples. Genes that showed significant signal of positive selection included *STP*1, *MAEBL, MSP*3.8, and *PIR* gene regions. *PvMSP*3.8 gene may play a role in the erythrocyte invasion. *MAEBL* is a membrane associated erythrocyte binding like protein that may have a function associated with erythrocyte invasion.

All *RBP* genes, except for *RBP*2c, showed regions with significant signals of positive selection (average d_N_/d_S_ ratio across all sites: 5.11; Table 2). Among them, *RBP2*p1 had the largest number of sites with posterior probabilities greater than 95% (Table 2). Among all the *MSP* genes, only *MSP*5, *MSP*9, and *MSP*10 indicated regions under positive selection. The *MSP*5 and *MSP*9 genes had an average d_N_/d_S_ ratio of 3.85 across all sites and 1.11 across branches (Table 2). While *MSP*10 had an average d_N_/d_S_ ratio of 1.16 across all branches and less than 1 across all sites, only seven sites were indicated with posterior probabilities greater than 50% and 95% (Table 2). Although *MSP*3.8 showed potential positive selection based on the integrated haplotype scores (Figure 4), PAML did not show significant evidence of positive selection. For the *DBP* gene family, *DBP*9 showed the highest d_N_/d_S_ ratio across all sites and branches (10.39 and 3.88, respectively; Table 2).

### Copy number variation and evolution of high-order copy variants

According to CNVnator, 19 gene regions showed copy number variation among our samples (Figure 5; Supplementary Table 4). Among them, 11 gene regions were detected with up to 2-3 copies and 8 gene regions with 4 copies or higher. We observed copy number variation in several *PIR* genes distributed across chromosomes 1, 2, 4, 5, 7, 10 and 12 (Figure 5; Supplementary Table 4). Specifically, for the *PIR* genes located on chromosome 2 (including PVP01_0220700, PVP01_0200200, PVP01_0200300, and PVP01_0200100; Figure 5), more than 20% of the samples had 2-3 copies and approximately 2-4% of the samples had 4 copies or higher. Among the 64 erythrocyte binding genes, duplications were observed in *DBP*1 on chromosome 6 and *MSP*3 on chromosome 10. *DBP*1 ranged from one to as high as five copies, and *MSP*3 ranged from one to as high as three copies among our samples (Figure 5), consistent with previous findings [19, 20, 76]. The remaining erythrocyte binding genes were detected with a single copy across our samples.

**Figure 5.**
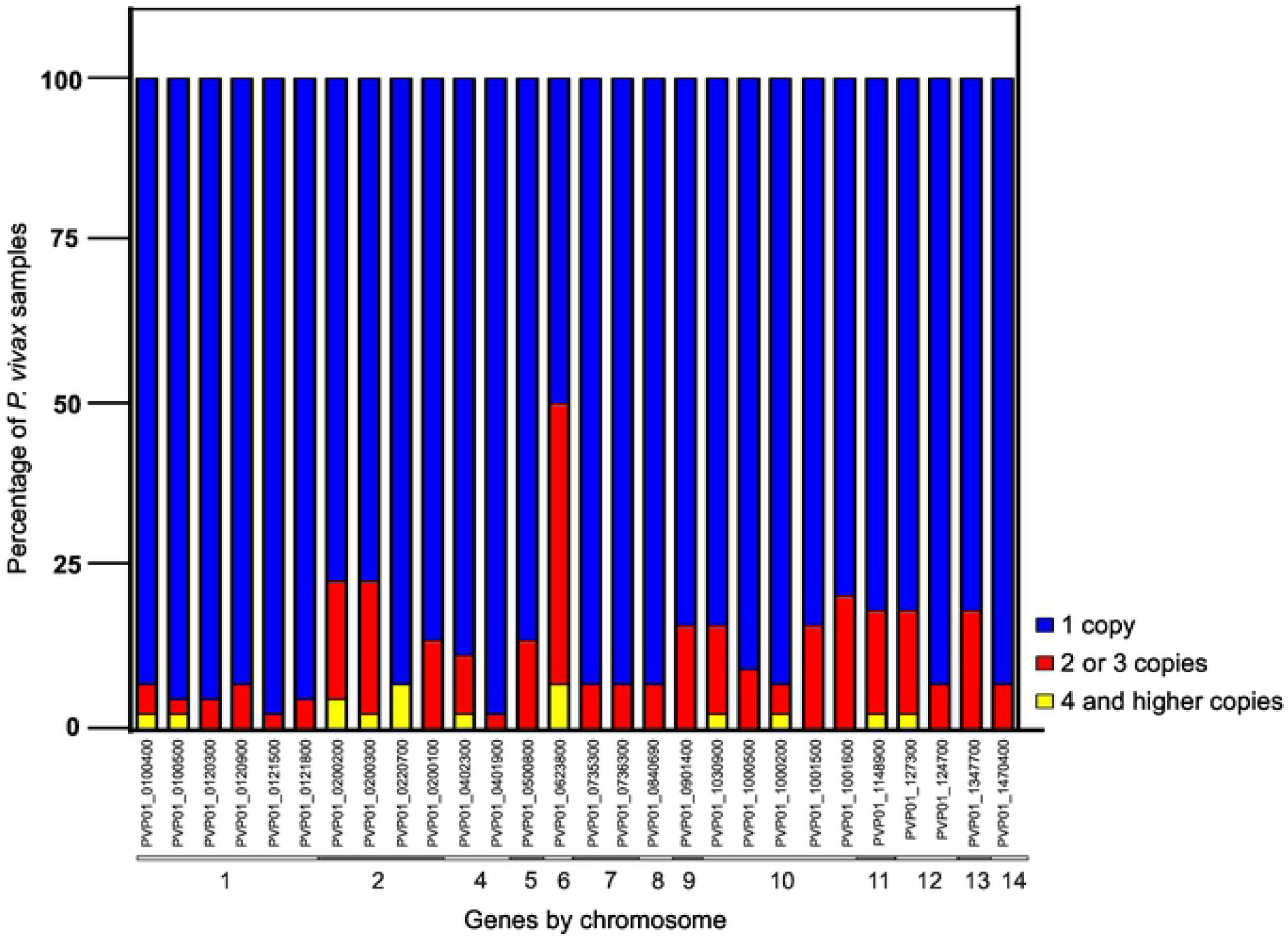
A total of 28 gene regions that were detected with copy number variation. Annotation of these genes can be found in Supplementary Table 4. Among them, *PvDBP*1 (PVP01_0623800) and *PvMSP*3 (PVP01_1030900) were associated with erythrocyte invasion. Other genes that were found to have high-order copy number were *PIR* protein genes or unknown exported plasmodium proteins.

A maximum likelihood tree constructed based on the whole genome sequences showed an admixture of *P. vivax* isolates with single and multiple *PvDBP* copy number (Figure 6A). The Ethiopian *P. vivax* isolates were divided into six subclades. Subclade I contained *P. vivax* samples mostly from Arbaminch and Badowacho with both one and two *PvDBP* copies. Subclade II contained samples from Jimma and Hawassa with two *PvDBP* copies. Subclade III contained a mixture of *P. vivax* samples from Arbaminch, Halaba, Hawassa, and Jimma with single and high-order *PvDBP* copies. This clade was sister to subclade IV that contained *P. vivax* samples mostly from Jimma (Figure 6A). In subclade IV, no distinct clusters were detected between isolates with single and multiple *PvDBP*. Subclade V contained samples from Jimma and subclade VI contained samples from Arbaminch, Badowacho, Hawassa, and Halaba. Each of the subclades had samples with both one and two *PvDBP* copies. Similar patterns were observed in the *MSP*3 and *PIR* genes where *P. vivax* isolates with single and multiple copies were clustered together in separate subclades (Figures 6B-D), suggesting that these gene regions could have expanded multiply among samples at different locations.

**Figure 6.**
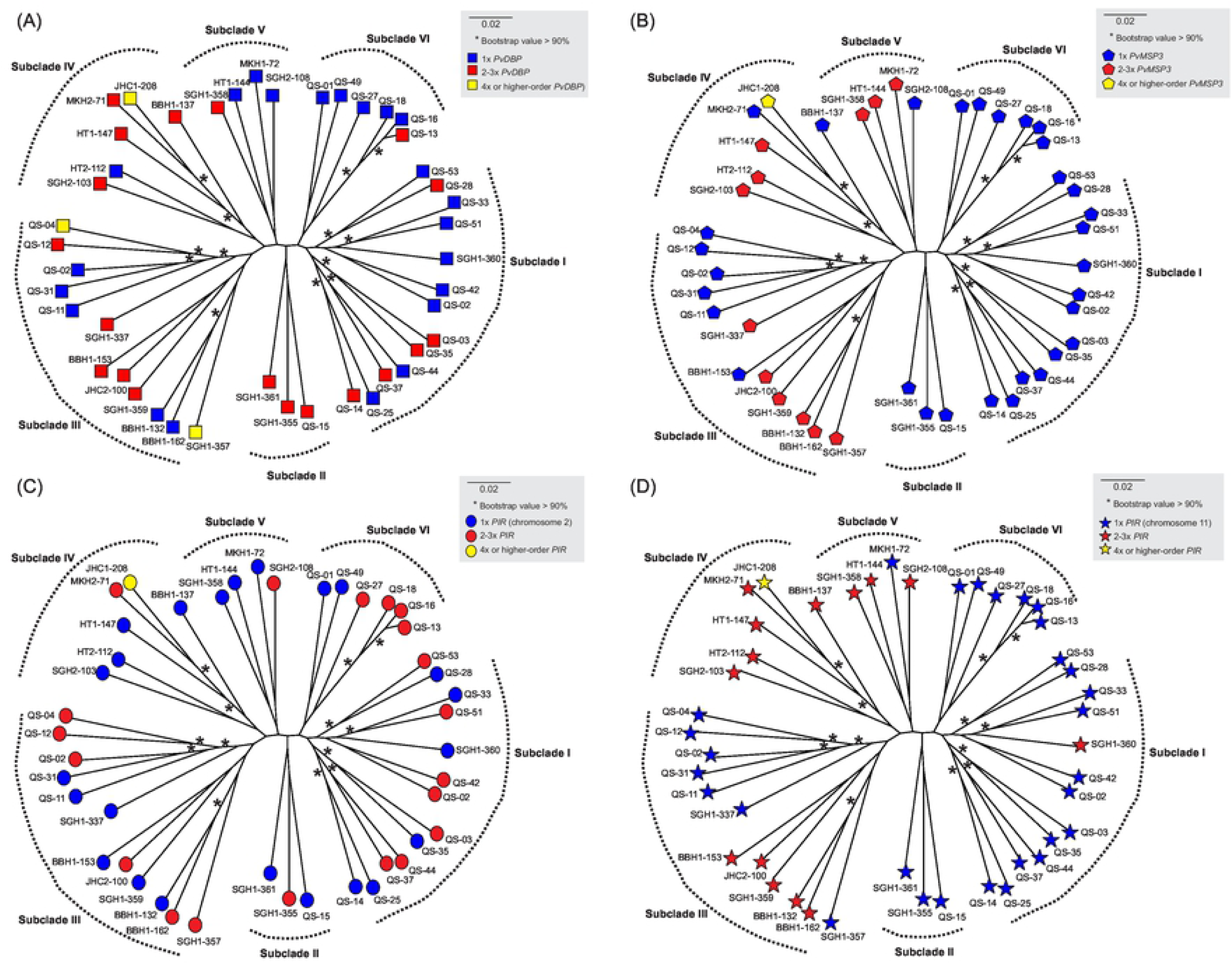
An unrooted whole genome phylogenetic tree of the 44 Ethiopian samples showing the evolution of (A) *PvDBP*; (B) *PvMSP*3; (C) *PIR* gene on chromosome 2; and (D) *PIR* gene on chromosome 11. The Ethiopian isolates were divided into three subclades. Subclade I contained samples mostly from the Arbaminch and Badowacho. Subclade II contained a mixture of isolates from Arbaminch, Halaba, Hawassa and Jimma. Subclade III contained samples from Jimma. No distinct clusters were observed between isolates with single and multiple *PvDBP, PvMSP*3, and *PIR* genes. These patterns suggest that these gene regions could have expanded multiply among samples at different locations.

### Gene flow and transmission network of the Ethiopian *P. vivax*

The principal component analysis based on the SNP variants showed samples from Arbaminch, Badowacho, Hawassa, and Halaba were genetically closely related but differentiated from Jimma (Figure 7A). The transmission network indicated that Arbaminch was the major source or hub of infections where the infections in Jimma, Hawassa, Badowacho, and Halaba were originated from (Table 3; Figure 7B). On the other hand, no transmission was originated from Halaba, making this location the largest sink of transmissions. The greatest extent of gene flow was observed between Arbaminch and Badowacho (Figure 7B). Hawassa and Jimma showed a source hub ratio of 0.5, indicating that there are equally as many egress transmissions as ingress transmissions (Table 3). Although Jimma and Badowacho/Halaba are in close geographical proximity, no apparent gene flow was observed between these sites.

**Figure 7.**
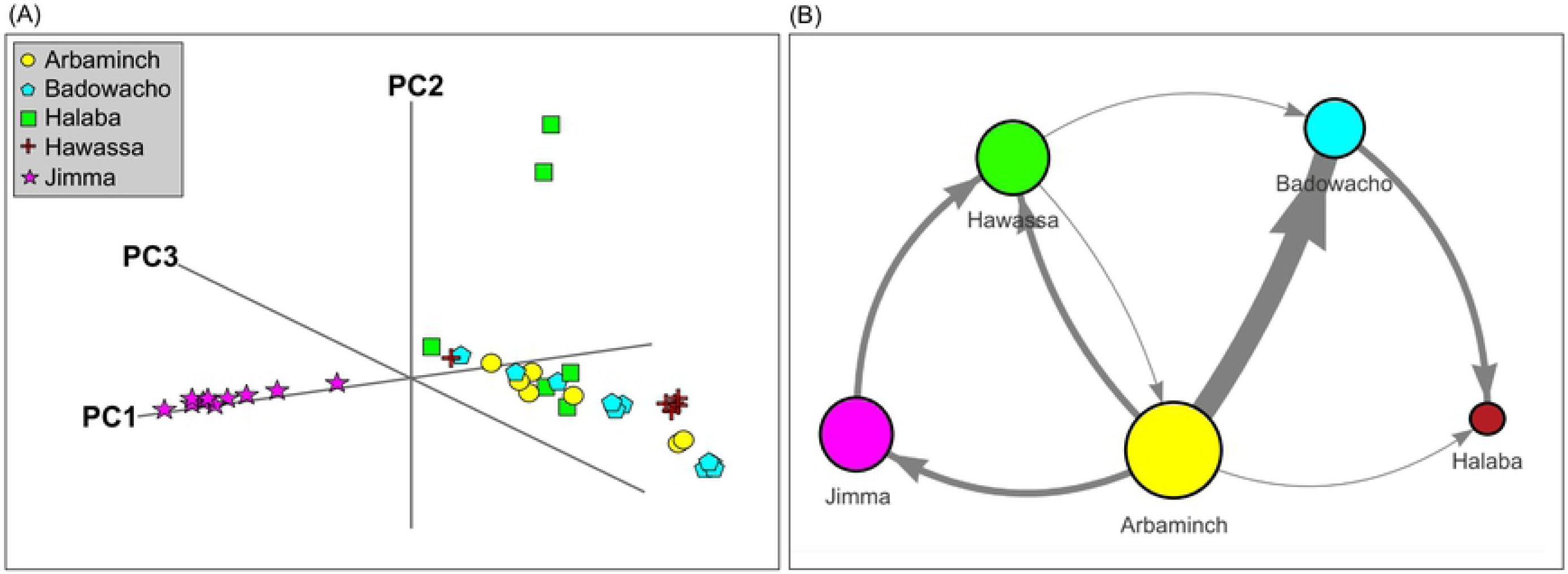
(A) Principal component analysis plot based on the SNP information from our variant analysis. Samples obtained from Jimma were clustered together, whereas samples from Arbaminch, Badowacho, Hawassa, and Halaba were mixed together with the exception of two samples from Hawassa. This clustering pattern suggested that there was considerable genetic variation among study sites even at a small geographical scale. (B) The transmission network, created using the StrainHub program, indicated that Arbaminch was the major source of infection in Jimma, Halaba, Badowacho and Hawassa. The greatest extent of gene flow (indicated by the boldest arrow) was observed between Arbaminch and Badowacho. Even though Jimma, Badowacho and Halaba are geographically in close proximity, gene flow was not intense among these sites.

## Discussion

Across the genome, the total number of SNPs observed among 44 *P. vivax* isolates in Ethiopia were comparable to those previously reported in South American [77] and Southeast Asian countries [19]. For instance, 303,616 high-quality SNPs were detected in 228 *P. vivax* isolates from Southeast Asia and Oceania in a previous study, of which Sal-I was used as the reference sequence and subtelomeric regions were discarded [19]. Auburn *et al*. [20] found that the average nucleotide diversity in Ethiopia was lower than in Thailand and Indonesia, but higher than in Malaysia. Chromosomes 3, 4, and 5 have been previously shown to contain the lowest proportion of synonymous SNPs than the other parts of the genome [12]. In the present study, chromosomes 3 and 6 were found to have the lowest number of both synonymous and nonsynonymous SNPs. This follows observations made in other studies done with nucleotide diversity ranging from 0.8 SNPs per kb in North Korea to 0.59 SNPs per kb in Peru [78]. Among the 64 erythrocyte binding gene candidates, the MSP and RBP multigene families showed the highest level of genetic variation. This agrees with previous studies that reported a remarkably high diversity in *RBP*2 than in *RBP*1 and its homolog group in *P. falciparum* [31]. In the Greater Mekong Subregion, the *MSP*3 and *PIR* gene families also indicated high levels of genetic diversity with 1.96% and 1.92% SNPs per base respectively, confirming that members of multigene families are highly variable genetically [30, 79]. Such diversity suggested that the binding domains of these genes could be under differential selection pressure. This pattern has been observed in previous studies and is likely due to their critical role in reticulocyte invasion, immunogenic properties, and human migration [26, 80-82].

Both CNVnator and GATK4 showed high order copies in several *PIR* gene regions. In addition, the *PIR* and *STP*1 genes were also indicated with significant selection based on the iHS calculations. The *PIR* gene family, which includes *STP*1, are located on the subtelomere regions and is a highly variable multigene family ranging from 1,200 genes in the reference strain PvP01 to 346 genes in monkey-adapted strain Salvador-I [56, 83]. Our analyses included only SNP variants that had a quality score of 40 or higher. Also, we used the PVP01 reference genome to map and annotate the subtelomeric regions, with the goal to reflect variability and features across the entire chromosome; whereas previous studies used the Sal-I reference genome with hypervariable and subtelomeric regions removed to minimize mapping errors [19, 84]. A recent study in *P. chabaudi* suggested that polymorphisms in *PIR* genes could affect the virulence of the parasites following passage from the mosquitoes [85]. Such a variation in copy number of the *PIR* gene family has also been reported in *P. cynomolgi* and *P. vivax* [86], suggesting that gene duplication could have been occurred repeatedly in the ancestral lineages [86]. The *PIR* multigene family is one of the largest gene families identified so far in *P. vivax* with several different potential functions. Some *PIR* genes encode proteins on the surface of infected red blood cells, which could confer to immune evasion; others encode proteins involved in signaling, trafficking and adhesion functions [83]. Positive selection detected in the *PIR* genes among the Ethiopian *P. vivax* isolates may have important implications on the susceptibility of the mosquito hosts [87].

For the *P. vivax* isolates in Southeast Asia, copy number variation was observed in nine gene regions including *DBP*1, *MDR*1, and PVX_101445 (on chromosome 14) with copy number ranging from 3 to 4 [19]. *DBP*1 and *MSP*3 showed higher order copies when compared to other genomic regions. In this study, the highest and most variable copy number variations were detected in the *DBP*1, with copy numbers ranging from one to as high as five. Likewise, for the *MSP*3, copy numbers ranging from one to as high as four. Based on the phylogeny, *DBP*1 and *MSP*3 expansion had occurred multiple times as tandem copies. These findings were consistent with earlier studies [19, 76] and suggested that gene expansion may play a key role in host cell invasion [88]. For all other putative erythrocyte binding genes, only a single copy was detected among all samples. A larger sample in future investigations would verify this observation.

In the present study, we identified a panel of 64 putative erythrocyte binding gene candidates based on the information from the literature and analyzed their polymorphisms. However, we did validate the function for each of these genes. Among these 64 putative erythrocyte binding gene candidates, *MAEBL* was shown to be highly conserved in *Plasmodium* [89], had the highest signal for positive selection among the *P. vivax* samples in Ethiopia. In *P. berghei, MAEBL* is a sporozoite attachment protein that plays a role in binding and infecting the mosquito salivary gland [89]. In *P. falciparum, MAEBL* is located in the rhoptries and on the surface of mature merozoites, and expresses at the beginning of schizogony [89]. In *P. vivax, MAEBL* is a conserved antigen expressed in blood stages, as well as in the mosquito midgut and salivary gland sporozoites [89, 90]. The *MAEBL* antigen contains at least 25 predicted B-cell epitopes that are likely to elicit antibody-dependent immune responses [91]. Positive selection observed in this gene region among the Ethiopian *P. vivax* isolates could be associated with the immunity-mediated selection pressure against blood-stage antigens. Though *DBP*1 had the highest and most diverse copy number variation, no significant signal of positive selection was detected.

It is noteworthy that the calculation of integrated haplotype scores and the accuracy of phasing genotypes using BEAGLE were dependent on the levels of linkage disequilibrium of the whole genomes. The higher the levels of linkage disequilibrium, the more accurate are the phased genotypes and thus the iHS score. Pearson *et al*. [19] found that *P. vivax* experienced drops in linkage disequilibrium after correcting for population structure and other confounders. Linkage disequilibrium of *P. vivax* genomes has been previously shown to be associated with the rate of genetic recombination and transmission intensity [92-94]. In high transmission sites of Papua New Guinea and the Solomon Islands, no identical haplotypes and no significant multilocus LD were observed, indicating limited inbreeding and random associations between alleles in the parasite populations [95, 96]. However, when transmission intensity declined, similar haplotypes and significant LD were observed possibly due to self-fertilization, inbreeding and/or recombination of similar parasite strains [92]. Multilocus LD is significantly associated with the genetic relatedness of the parasite strains [97], but inversely associated with the proportion of polyclonal infections [98]. In Southwestern Ethiopia, malaria transmission ranged from low to moderate, and LD levels varied markedly among the study sites [53, 99]. To address this limitation in BEAGLE, all genes that were detected with positive selection in BEAGLE were further analyzed with PAML for verification. Future study should include broad samples to thoroughly investigate selection pressure at the population level and the function significance of polymorphisms in the *MAEB*L and *PIR* genes.

Previous studies have shown high levels of genetic diversity among *P. vivax* isolates in endemic countries [16, 100, 101]. Such a diversity was directly related to high transmission intensity and/or frequent gene exchange between parasite populations via human movement [4, 12, 13, 53]. For example, previous studies using microsatellites have demonstrated a consistently high level of intra-population diversity (*H*_E_ = 0.83) but low between-population differentiation (*F*_ST_ ranged from 0.001-0.1] in broader regions of Ethiopia [53, 99]. High heterozygosity was also observed in *P. vivax* populations from Qatar, India, and Sudan (average *H*_E=_ 0.78; 62), with only slight differentiation from *P. vivax* in Ethiopia (*F*_ST_ = 0.19) [102]. Frequent inbreeding among dominant clones [92, 95] and strong selective pressures especially in relapse infections [19, 20, 102, 103] may also contribute to close genetic relatedness between and within populations. Thus, in this study, it is not surprising to detect a high level of parasite gene flow among the study sites at a small geographical scale, despite the limited number of samples. In the present study, we successfully employed a transmission network model to identify transmission paths, as well as the source and sink of infections in the region, beyond simply indicating genetic relationships.

To conclude, this study elaborated on the genomic features of *P. vivax* in Ethiopia, particularly focusing polymorphisms in erythrocyte binding genes that potentially play a key role in local parasite invasion, a critical question given the mixed Duffy positive and negative populations of Ethiopia. The findings provided baseline information on the genomic variability of *P. vivax* infections in Ethiopia and allowed us to compare the genomic variants of *P. vivax* between Duffy-positive and Duffy-negative individuals as the next step of our ongoing investigation. Further, we are in progress of developing a panel of informative SNP markers to track transmission at a micro-geographical scale.

## Data Availability

Additional information is provided as supplementary data accompanies this paper. Sequence data of this study are deposited in the European Nucleotide Archive (ENA) and the accession number of each sample is listed in Table 1.

## Acknowledgements

We are greatly indebted to the staffs and technicians from Jimma University for field sample collection, the communities and hospitals for their support and willingness to participate in this research.

## Tables

**Table 1.** Information of whole genome sequences of 44 *Plasmodium vivax* isolates from Ethiopia. The European Nucleotide Archive (ENA) accession number for all files.

**Table 2.** A shortlist of 25 erythrocyte binding genes that showed signals of positive selection based on the Likelihood Ratio Test of the M1 (neutral model) and M2 models (selection model) in PAML.

**Table 3.** Transmission network metrics among study sites calculated by StrainHub.

## Supplementary files

**Supplementary Table 1.** Distribution of SNP variants in the 64 *P. vivax* erythrocyte binding gene candidates among the 44 Ethiopian genomes.

**Supplementary Table 2.** Distribution of single nucleotide polymorphism (SNP) variants across *P. vivax* chromosomes of the 44 Ethiopian genomes.

**Supplementary Table 3.** Likelihood Ratio Test results of the M1 (neutral model) and M2 models (selection model) in PAML of all the 64 erythrocyte binding gene candidates.

**Supplementary Table 4.** Gene regions that were detected with copy number variation among the 44 Ethiopian *P. vivax* isolates based on CNVnator. Among them, only two erythrocyte binding gene candidates *PvDBP*1 and *PvMSP*3 were detected with high-order copies.

